# Virus coinfection in plants determined by ecological responses to environmental heterogeneity

**DOI:** 10.64898/2026.11.04.764184

**Authors:** Adrian D. Zamfir, Bisola M. Babalola, Michael McLeish, Fernando García-Arenal

**Author notes:** **Corresponding author**: Fernando García-Arenal, Mailing address: CBGP, Campus de Montegancedo, UPM, 28223 Pozuelo de Alarcón, Madrid, Spain.

## Abstract

Community ecology is acknowledged as a relevant causative agent of epidemiological processes of plant viruses and has consequences for coinfection and virus and host evolution. Plant virus coinfection is common, but factors that promote non-random virus interactions above the population-level are unknown. Here, we analyse intrinsic and extrinsic factors in virus-host and virus-virus interactions by eighteen generalist virus species of a diverse multi-host – multi-virus system. Coinfection among viruses with broad host ranges, diverse modes of horizontal transmission, and interspecific variation in responses to environmental heterogeneity is expected to be shaped by ecological as well as evolutionary constraints. We analyse patterns of coinfection from 1,931 individual plants in 94 species from multiple plant communities of distinct habitat types and show that above 50% of plants were coinfected by 2-8 viruses. To analyse the role of environmental variables on the observed virus associations we used Markov and conditional random fields modelling. Results showed that a large majority of coinfections were random interactions, consistent with ecological fitting, and only a minor fraction being non-random associations. Random field models underscored the relevance of common responses to environmental heterogeneity driving coinfection, and strongly suggest mode of horizontal transmission as a causal agent. Results highlighted a major role of between-host, rather than within-host processes in driving coinfection. This study is a considerable addition to the understanding of drivers of virus coinfection and, hence, of virus community ecology and epidemiology.

## INTRODUCTION

In nature, individual hosts are often coinfected by various parasites, which may have consequences for epidemiology and parasite and host evolution. Theoretical studies of parasite coinfection (i.e., more than a single parasite detected from an individual host) has a long history, however, further empirical support is necessary (1) (2–6). Coinfection by plant viruses has received much attention (7, 8) since an early finding that the “rugose mosaic” disease of potato is caused by an interaction between potato virus X and potato virus Y (9), which lead to the discovery of synergistic interactions between coinfecting viruses (10). A wealth of subsequent experimental studies has shown that virus interactions in coinfected plants can be neutral, antagonistic or facilitative to one or both partners. Recent reports indicate that plant virus coinfection is common (11–16) (17, 18), but little is known about the mechanisms that promote coinfection. Importantly, coinfection is expected to be shaped by non-random associations determined by mechanisms intrinsic to virus-host or virus-virus interactions, as well as factors extrinsic to that association.

The community ecology of plant viruses in nature (18, 19) has recently been acknowledged as a causative agent important to epidemiological processes. The mechanisms underlying coinfection have been explained by complementation or competition between virus-encoded functions, and/or to manipulation of host defences (7, 8, 20, 21). It has also been shown that coinfection can affect the efficiency of virus transmission by vectors (22) via altered virus multiplication, facilitative or antagonistic virus interactions in the vector itself (23, 24), and the manipulation of vector behaviour (25). Knowledge of the effects of coinfection on plants and vectors suggest virus intrinsic factors that act within and between hosts are major determinants of non-random virus associations. Intrinsic factors are related to common evolutionary histories that condition infection, replication, and transmission pathways (4). However, non-random associations can also result from extrinsic factors (26, 27). For instance, it was shown that virus-virus associations depended on the spatial distributions of host plant species (13, 28, 29). Similar trait responses of different viruses to environmental heterogeneity, and interaction of otherwise independent viruses with a common virus partner, have also been highlighted as extrinsic factors in non-random associations (4). To our knowledge, drivers of non-random associations among viruses infecting wild hosts have only been conducted at the plant population level (18) and not at the community scale.

Here, we investigate and test patterns of coinfection by eighteen generalist virus species that have broad host ranges across hosts from multiple plant communities (30, 31). The 18 viruses belong to nine genera and represent different modes of horizontal transmission and diverse evolutionary histories. Infection was detected from RNA extracts of individual plants by RT-PCR with virus-specific primers, appropriate for the detection of coinfection (we make no distinction with superinfection). We use Markov and conditional random fields modelling (26) to analyse the role of biotic and abiotic environmental variables on the observed virus associations. We estimate the prevalence of coinfection and address the following null hypotheses: i) the proportion of random and non-random associations is equivocal; and (ii) extrinsic factors have no effect on the proportion of random to non-random associations.

## MATERIAL AND METHODS

### Study area and plant sampling

Plants were sampled at field sites between July 2015 and June 2017 in the Vega del Tajo-Tajuña agricultural region in the South-Central plateau of the Iberian Peninsula (Spain) (**Figure S1**). Repeat sampling was conducted at multiple sites of each of four habitats (**Table S1**) that were subject to different degrees of disturbance. Oak forests (Oak) are native communities dominated by evergreen *Quercus* spp.; Wasteland includes undisturbed patchy areas of successional shrubland with no human use for the last ten years; Crop are habitats where annual monocultures share the space with communities of wild plant species (weeds) that reassemble every year, and Edge are the narrow borders that separate crop fields, with plant communities that benefit from nutrient and water supplemented to adjacent crops. Oak and Wasteland are considered as non-anthropic habitats, with presently undisturbed communities, while Crop and Edge together formed the anthropic habitats. Seventy-eight collections were performed at four sites each for Oak, Wasteland and Edge habitats, and eleven sites of Crop.

The higher number of Crop sites aims to represent the variation expected from the Crop communities as fields are cultivated with different summer (melon and maize) and winter (barley and brassicas) crop species, and are subject to crop rotation and fallow periods. Collections at sites of Oak and Wasteland were conducted in spring and autumn, at sites of Crop when the summer or winter crops were present, and at sites of Edge in spring and autumn and when the adjacent crops were sampled. Plants were sampled systematically along fixed itineraries, regardless of whether or not they showed symptoms of diseases or herbivory. The sampling design (described in detail in (30) maintained the relative species richness and abundance of plant species distributed among the four habitats. Previous work has demonstrated differences in plant species diversity among the communities that characterise each habitat, with positive spatial autocorrelation at broad scales (i.e., ecosystem) driven by the environment (30). Significant negative spatial autocorrelation at fine-scales showed community structuring among adjacent study sites of the different habitat categories. As study sites were generally not contiguous, with intervening patches surrounding the sites, explicit consideration of spatial scale, geographic distance, and connectivity measures between sites, critical to the interpretation of biological processes, was given precedence over habitat area (30).

### Virus detection

Total RNA from individual plant species was extracted with the Agilent Plant RNA Isolation Mini Kit, Spectrum^TM^ Plant Total RNA Kit, and the CTAB method (32) depending on the plant species. To minimise contamination across samples, extractions were conducted in groups of samples from the same species and collection (i.e. same time, same site). Plant tissues were ground in liquid nitrogen with mortar and pestle (one for each individual sample) before adding the extraction reactants. Mortars and pestles were washed in 0.2 M NaOH and autoclaved before being used. RNA extracts from same species and collection were pooled to obtain a single high throughput sequencing (HTS) library. Both read orientations of each pool were run in the same sequencing lane, and each pool run in a separate lane. All libraries included a step for rRNA depletion using the RiboZero kit. Library preparation and high throughput sequencing with Illumina HiSeq platforms were as in (33). This yielded 323 libraries of 2,037 pooled individual samples representing 119 plant species that were sampled at least 5 times in a single habitat.

In these libraries a total of 158 virus OTUs had been detected (34), of which 18 OTUs that were detected in hosts that occurred in more than one habitat and that have diverse life histories were selected for the present study. The 18 focal OTUs matched references of virus species of 9 genera with positive-sense single-stranded RNA genomes, and were transmitted horizontally by aphids in a non-persistent or a persistent non-propagative manner, by pollen, by contact or by Coleoptera. These virus species differed in genome structure and evolutionary history as represented across the 9 genera. In some instances, species of different genera shared a mode of horizontal transmission. The 18 virus species were: Pelargonium zonate spot virus (*Anulavirus PZSV,* PZSV), turnip crinkle virus (*Betacarmovirus brassicae,* TCV), cucumber mosaic virus (*CucumovirusC MV,* CMV), tomato aspermy virus (*Cucumovirus TAV,* TAV), parietaria mottle virus (*Ilarvirus PMV,* PMoV), barley yellow dwarf virus-PAV (*Luteovirus mavhordei,* BYDV-PAV), beet chlorosis virus (*Polerovirus BCHV,* BChV), beet mild yellowing virus (*Polerovirus BMYV,* BMYV), cucurbit aphid-borne yellows virus (*Polerovirus CABYV,* CABYV), turnip yellows virus (*Polerovirus TUMV,* TuYV), plum pox virus (*Potyvirus plumpoxi,* PPV), turnip mosaic virus (*Potyvirus rapae,* TuMV), watermelon mosaic virus (*Potyvirus citrulli,* WMV), rubus chlorotic mottle virus (*Sobemovirus SOMV,* RuCMV), pepper mild mottle virus (*Tobamovirus capsici,* PMMoV), tobacco mild green mosaic virus (*Tobamovirus mititesselatti,* TMGMV), tobacco mosaic virus (*Tobamovirus tabaci,* TMV) and Youcai mosaic virus (*Tobamovirus youcai,* YoMV). Information on species taxonomy and mode of transmission is given in **Table 1**. The occurrence in individual plants of the 18 viruses was analysed by RT-PCR using sets of specific primers that amplified regions of 70–174 nucleotides in the coat protein gene, as in (31) (**Table S2**). Primer sequences were a consensus of the sequences for that virus species in the available databases. For RT-PCR, the complementary DNA (cDNA) was synthesized with Maxima First Strand cDNA Synthesis Kit (ThermoFisher Scientific, Lithuania) using 0.5 µg of individual plant RNA extracts. The PCR was performed with Supreme NZY Taq II polymerase kit (NZYtech, Lisboa, Portugal) with the following PCR cycle conditions: initial denaturation at 95°C for 3 minutes, 40 cycles of denaturation at 94°C for 30 seconds, annealing temperature based on each primer pair for 30 seconds, and extension at 72°C for 30 seconds. Lastly, a final extension of 72°C for 7 minutes was carried out. The PCR products were visualised by gel electrophoresis in 2.0% agarose after GelRed^®^ staining. RT-PCR with virus-specific primers was performed on a subset of randomly selected libraries in which OTUs of the 18 focal viruses were detected by HTS (34), on the condition that at least 50% of the libraries of each habitat were represented.

**Table 1.** Taxonomy and mode of transmission of eighteen virus detected by RT-PCR in individual plant samples.

| **Virus Genus** | **Virus species** | **Mode of horizontal transmission** |
|---|---|---|
| ***Anulavirus*** | PZSV | Pollen |
| ***Betacarmovirus*** | TCV | Contact, coleoptera |
| ***Cucumovirus*** | CMV | Aphid, non-persistent |
| ***Cucumovirus*** | TAV | Aphid, non-persistent |
| ***Ilarvirus*** | PMoV | Pollen |
| ***Luteovirus*** | BYDV | Aphid, persistent non propagative |
| ***Polerovirus*** | BChV | Aphid, persistent non propagative |
| ***Polerovirus*** | BMYV | Aphid, persistent non propagative |
| ***Polerovirus*** | CABYV | Aphid, persistent non propagative |
| ***Polerovirus*** | TuYV | Aphid, persistent non propagative |
| ***Potyvirus*** | PPV | Aphid, non-persistent |
| ***Potyvirus*** | TuMV | Aphid, non-persistent |
| ***Potyvirus*** | WMV | Aphid, non-persistent |
| ***Sobemovirus*** | RuCMV | Contact |
| ***Tobamovirus*** | PMMoV | Contact |
| ***Tobamovirus*** | TMGMV | Contact |
| ***Tobamovirus*** | TMV | Contact |
| ***Tobamovirus*** | YoMV | Contact |

### Analyses of virus associations in coinfected plants

The above detection procedure produced a data set of 1013 individual plants infected by at least one of the 18 analysed viruses, distributed across 23 sampling sites and 94 host plant species, generating a matrix of 18 columns (viruses) and 292 rows (host plant species x site) (**Table S3**). We detected non-random virus associations and the role of extrinsic factors on the co-occurrence of two viruses in coinfected plants, using Markov Random Fields (MRFs) models that estimate interspecific interactions (26, 35), and Conditional Random Fields (CRFs) models that allow for conditional dependencies on other environmental variables (36, 37). The models consist of networks comprised of nodes (virus species and environmental covariates/variables) and edges (the probabilistic effects of environmental variables on virus species occurrences plus the direct and indirect interactions occurring between the viruses), where edge effects are undirected and effect sizes between pairs of nodes are reciprocal (35). If there is no edge between two nodes in the network, these nodes are conditionally independent from one another, and if there is an edge, these nodes are conditionally dependent, after accounting for the other node effects in the network. The values for edge associations in the networks vary depending on the inclusion of species and environmental variables. Pairwise interactions were modelled as in (35). The log-odds of observing virus species *i* given the presence/absence of virus species *j* and covariate *x* is:

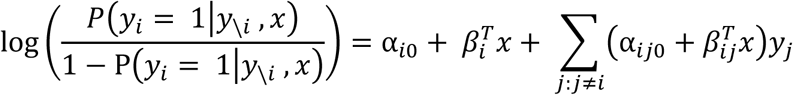

where *y_i_* is the vector of binary observations (presences/absences) of virus species *i*; y*_\i_* is a vector of binary observations of all other virus species except *i*; α*_i0_* is the virus-level intercept; and *β_i_^T^x* is the effect of covariate *x* on the occurrence probability of virus species *i*. Parameters *α*_ij0_ and *β_ij_^T^* represent the associations between virus species *i* and *j*, conditional on the occurrences of all the other viruses. The R package ‘MRFcov’ (v1.0.38) was used to perform these computations.

A series of environmental variables related to the spatial distribution among study sites, underlying spatial dependencies among communities, climate, habitat, and host plant traits, that were used in the random field analyses are listed in **Table 2**. The coordinates of the 23 field sampling sites were used to capture spatial structures using Moran’s Eigenvector Maps (MEMs) as in (30). The MEMs that best indicated spatial structuring (i.e., spatially explicit variation in species relative abundances among the communities) at the broad (MEM2, MEM7) and fine (MEM21) spatial scales. Gabriel graphs were used to describe distance relationships among sites (30) that were used in the ordination technique for generating the MEM’s.

**Table 2.**
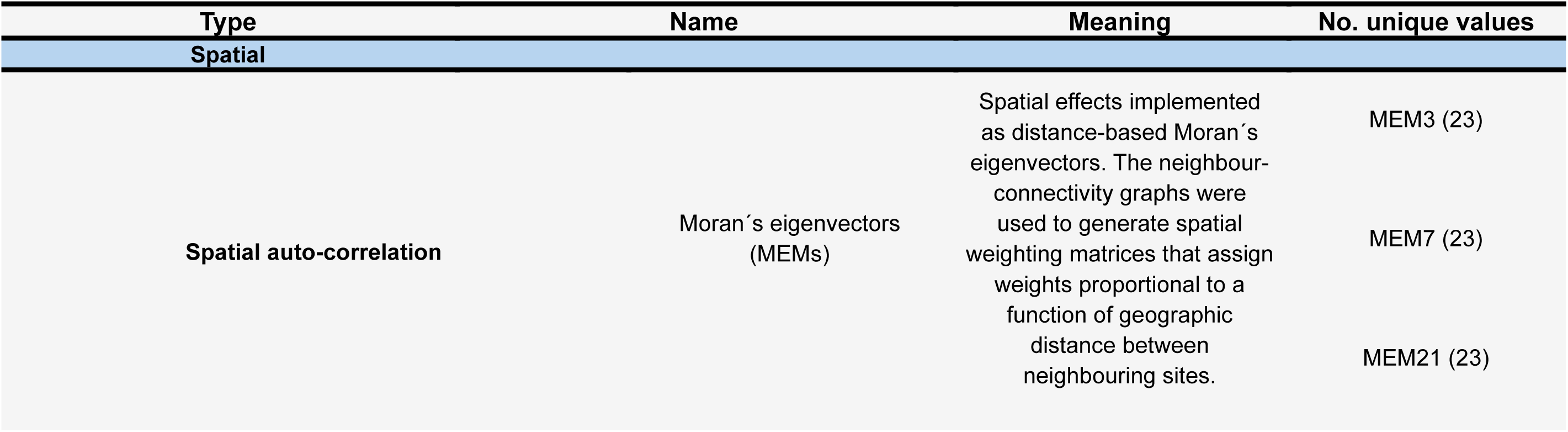
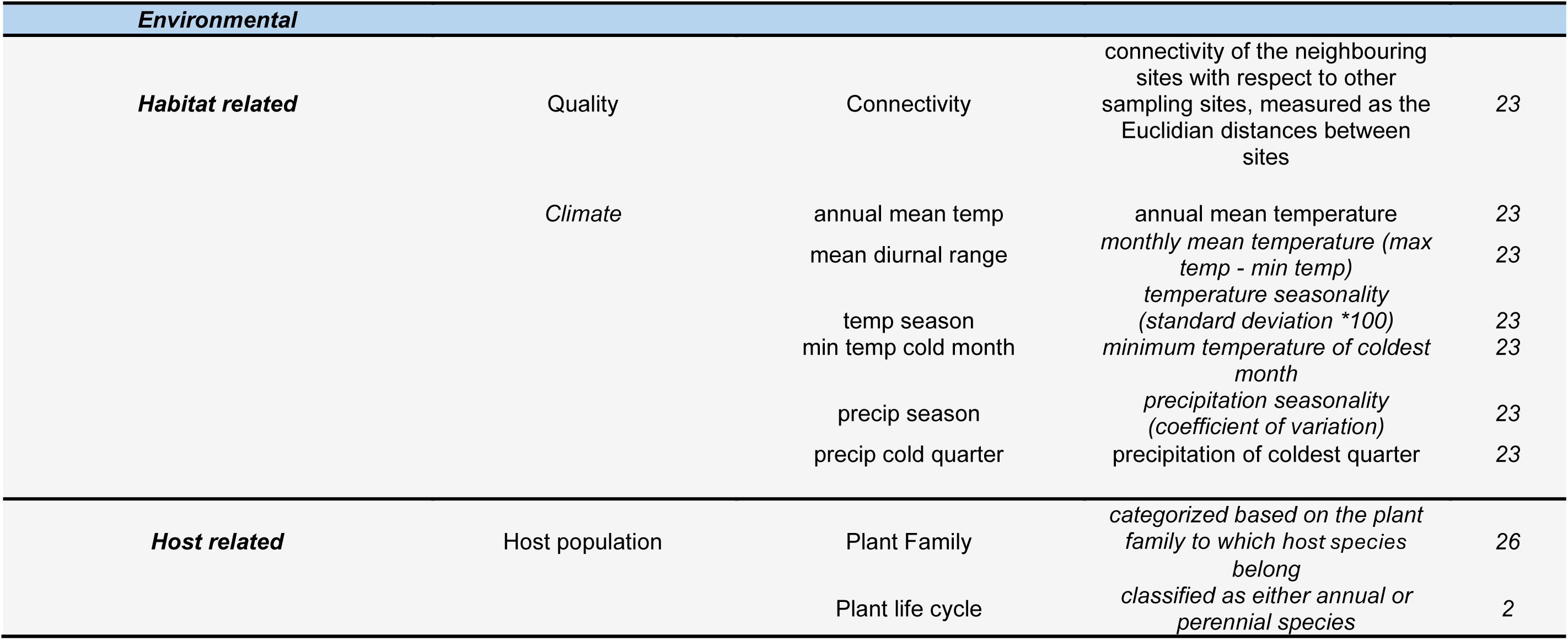
Spatial and environmental variables used for the CRF (conditional random field) models.

| Type |  | Name | Meaning | No. unique values |
|---|---|---|---|---|
| **Spatial** |  |  |  |  |
| **Spatial auto-correlation** |  | Moran´s eigenvectors (MEMs) | Spatial effects implemented as distance-based Moran´s eigenvectors. The neighbour-connectivity graphs were used to generate spatial weighting matrices that assign weights proportional to a function of geographic distance between neighbouring sites. | MEM3 (23) |
|  |  |  |  | MEM7 (23) |
|  |  |  |  | MEM21 (23) |
| ***Environmental*** |  |  |  |  |
| ***Habitat related*** | Quality | Connectivity | connectivity of the neighbouring sites with respect to other sampling sites, measured as the Euclidian distances between sites | *23* |
|  | *Climate* | annual mean temp | annual mean temperature | *23* |
|  |  | mean diurnal range | *monthly mean temperature (max temp - min temp)* | *23* |
|  |  | temp season | *temperature seasonality (standard deviation \*100)* | *23* |
|  |  | min temp cold month | *minimum temperature of coldest month* | *23* |
|  |  | precip season | *precipitation seasonality (coefficient of variation)* | *23* |
|  |  | precip cold quarter | precipitation of coldest quarter | *23* |
| ***Host related*** | Host population | Plant Family | *categorized based on the plant family to which host species belong* | *26* |
|  |  | Plant life cycle | *classified as either annual or perennial species* | *2* |

Habitat-related variables included the connectivity between pairs of sites and six climate predictor variables (**Table 2**) that our previous work had shown had significant effects on dependencies among sampling sites (30). Last, we included two categorical variables related to the host plant population, the taxonomy at the family level, and the life cycle (annual or perennial) of the plant species. The environmental variables were incorporated into the model either all together or in subsets, so that different CRF models were considered: the CRF-Full model, with all the variables in **Table 2**, and models with only the spatial (CRF-MEMs), environmental (CRF-Environmental), habitat (CRF-Habitat) or host (CRF-Host) covariates. The dimensions and number of variables of the matrices for the MRF and CRF models are shown in **Table S4**, and the matrices for the models are shown in **Tables S5 to S11**.

The performance of the different models was assessed by means of Receiver operating characteristic (ROC) curves that compare the sensitivity (the proportion of correctly classified positive observations) and specificity (the proportion of correctly classified negative observations) of each model classifiers (38). We used the area under the ROC curve (AUC) to quantify the model’s ability to classify observations. The analysis was implemented in the R package ‘pROC’ (v1.18.5 (39). The mean predictive performance of each model for all virus species simultaneously was compared by 10-fold cross-validation based on the following parameters: Mean positive predictions, Mean total predictions, Mean sensitivity and Mean specificity. We generated 100 bootstrapped replicates for each model. If the 90% confidence interval of bootstrapped coefficients were distinct from zero, we considered the variable to have a statistically significant effect, following (18).

## RESULTS

### Prevalence and distribution of coinfection across plant communities

A total of 4,170 RT-PCR reactions were performed from RNA extracts of 1,931 individual plant samples that represent 94 host species. Of these, 1013 were infected by at least one virus. Only one virus was detected in 454 (44.8%) samples and more than one and up to eight different viruses in 559 (55.2%) samples that were coinfected (**Figure 1A**). Of the 1013 infected plants, 12.7% were infected by four or more viruses (**Table S12**).

**Figure 1.**
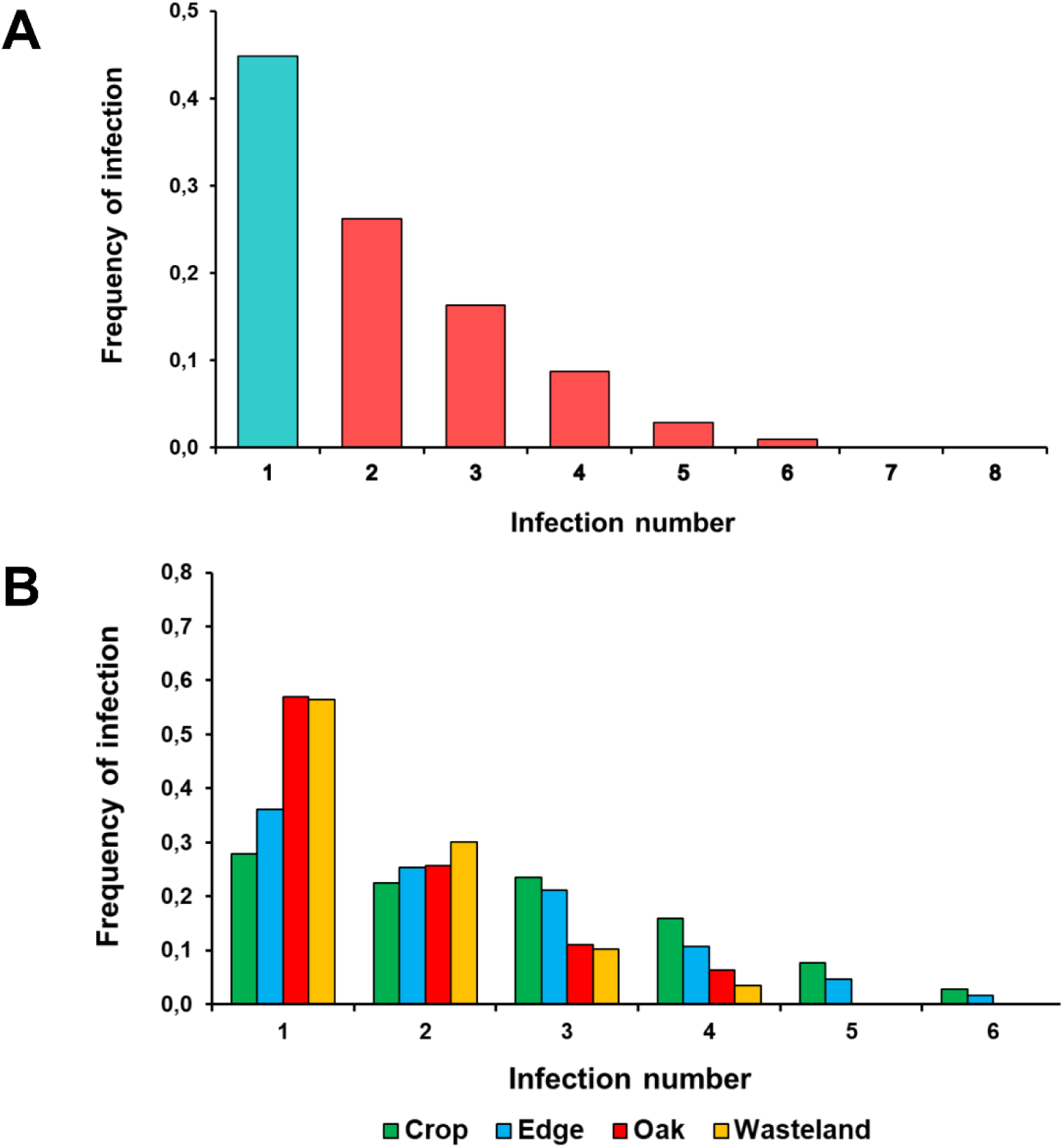
Distribution of the number of viruses infecting a plant at the ecosystem scale (A) and per habitat (B). Infection number indicates number of viruses detected per plant. Alt text: Distribution of virus coinfections per individual plant (A) or across habitats (B).

The prevalence of coinfection differed across habitats (χ²_3_ = 59.29, *p* < 0.001), with the frequency of coinfected plants being higher in Crop and Edge than in Oak and Wasteland (χ²_1_ >22.80, *p* < 0.001), with no difference between Crop and Edge (χ²_1_ = 3.39, *p* = 0.07) or between Oakwood and Wasteland (χ²_1_ = 0.002, *p* = 0.96). Also, the distribution of the number of viruses detected per individual plant differed among habitats (*χ*^2^_15_ = 154.18, *p* < 0.001) (**Figure 1B** and **Table S13**), again differing significantly between Crop and Edge vs. Oak and Wasteland (χ²_5_ > 39.15, *p* < 0.001), but not between Crop and Edge or between Oak and Wasteland χ²_5_ < 8.39, *p* > 0.968). The anthropic habitats Crop and Edge had more coinfected plants, and by a larger number (up to 8 in Edge) of virus species, than the non-anthropic habitats Oak and Wasteland, in which a maximum of four viruses were detected per individual plant. Thus, the pattern of coinfection varied according to the habitat, which suggests a role of extrinsic factors in coinfection.

### Extrinsic factors in coinfection

To test whether extrinsic factors had a role in determining coinfection, and in non-random virus-virus associations in coinfected plants, we used random field approaches to estimate the probability of two viruses being found in coinfection. Markov Random Field (MRF) networks were used to control for indirect effects of multispecies virus interactions, and Conditional Random Field (CRF) networks were used to evaluate conditional dependencies on environmental variation. The performance of the different models was evaluated using the Area Under the Curve (AUC) scores for their respective Receiver Operating Characteristic (ROC) curves. AUC scores showed that the addition of conditional environmental and spatial variables did not improve the performance of the MRF model (AUC = 0.887). The performance of the CRF models that included the spatial (CRF-MEMs, AUC = 0.883) and host-related covariables (CRF-Host, AUC = 0.881) was similar to that of the MRF model. However, the performance of CRF-MEMs and CRF-Host was significantly lower than that of MRF for one and three virus species, respectively (**Table S14**). AUC values for CRF-Full, CRF-Habitat and CRF-Environmental were lower (0.772, 0.741 and 0.719, respectively), and the performance of these three models was significantly lower than that of MRF for 11, 10 and 12 viruses, respectively (**Table S14**). Thus, the MRF, CRF-MEMs, and CRF-Host models were chosen for cross validation. Cross-validation indicated that the performance of CRF-MEMs and CRF-Host was close to that of MRF, values of the predictive performance parameters being similar at the different bootstrap quantiles (**Table S15**), while they were lower for the other models, which showed lower capacity to predict true positives (shown for CRF-Habitat in **Table S15**).

The MRF model (**Figure 2A**) yielded fifteen significant non-random interactions, with association coefficients between nodes at the 90% bootstrap confidence interval (see **Figure S2** for summary statistics). Two of these interactions were negative (TCV - PZSV; and TuYV - WMV) and the rest positive. Seven interactions were between viruses with the same mode of transmission. There were strong positive interactions between three aphid persistently transmitted poleroviruses (BChV, BMYV and TuYV), the two pollen transmitted viruses (PMoV and PZSV). There were weak positive interactions between two tobamoviruses (PMMoV and TMGMV) and TCV-YoMV, all contact transmitted, and between CMV and TAV and WMV that are aphid non-persistently transmitted.

**Figure 2.**
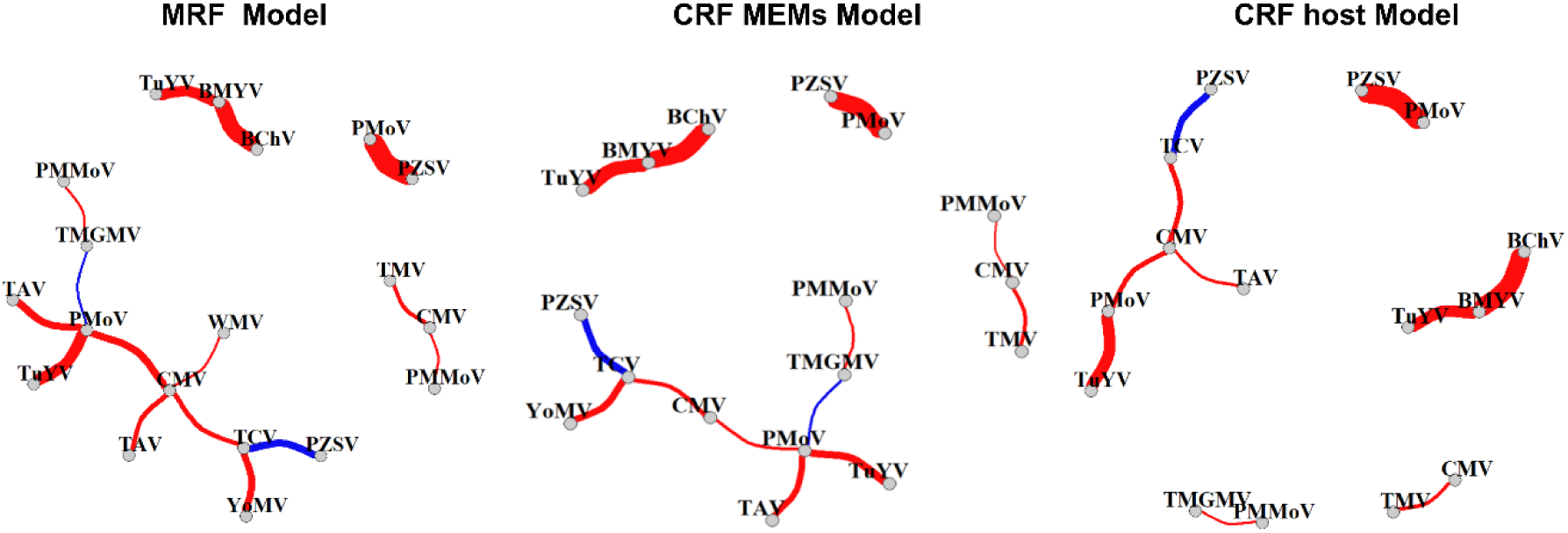
Virus interaction networks. A) Markov random field interaction network. B) Conditional Markov random field interaction network accounting for spatial variables (MEMs). C) Conditional Markov random field interaction network accounting for the host variables**. I**nteractions with an absolute edge weight ≤ 0.05 are not shown in the networks. Alt text. Associations between pairs of viruses retrieved by different random field models.

In addition, the MRF network revealed a set of virus interactions that were not structured according to a shared evolutionary history and/or mode of transmission. Here, CMV formed a hub, being the most connected virus, with PMoV, TCV, PMMoV and TMV in addition to TAV and WMV. The CRF-MEMs model (**Figure 2B**) retrieved thirteen significant interactions at the 90% bootstrap confidence interval (see **Figure S3** for summary statistics), two of them negative (PZSV – TCV as in MRF, and TMGMV – PMoV). The CRF-Host model (**Figure 2C**) retrieved ten significant interactions at the 90% bootstrap confidence interval (see **Figure S4** for summary statistics), all of them positive. The positive interactions of the CRF-MEMs and CRF-Host models included viruses belonging to the same genus (BChV, BMYV, TuYV and PMMoV, TMGMV), and between the pollen transmitted viruses (PZSV and PMoV) that the MRF model revealed. The interactions among viruses with different modes of transmission or belonging to different genera retrieved by the CRF-Host model, comprised a subset of those retrieved by the CRF-MEMs model, which were a subset of those shown by the MRF model. Spatial factors did not play a role in the interactions between CMV and TAV, which were retrieved in the MRF and CRF-Host models, but not in the CRF-MEMs. Host factors did not play a role in the interactions between TAV and PMoV, nor between TCV and YoMV, which were retrieved in the MRF and CRF-MEMs models, but not the CRF-Host model.

To disentangle and test the effects of the environmental covariables (**Table 2**) on virus community structuring, networks were generated in which the adjacency matrices of the CRF models that included a set of variables (**Table S4**) were subtracted from the adjacency matrix of models that included a larger set of the variables in **Table 2** (also including the MRF adjacency matrix). The network from the resulting matrix indicates the interactions are due to the variables not subtracted. Thus, the full environmental network (**Figure 3A**) was generated from the subtraction of the MRF matrix from that of the CRF-Full model, with all spatial and environmental variables. This network, based on the spatial and environmental covariates, revealed a core set of interactions that were associated with shared evolutionary history and/or mode of transmission (BChV, BMYV and TuYV; PMoV and PZSV; PMMoV and TMGMV). In addition, CMV and PMoV were hubs in interactiions with PMMoV, TCV, TAV, WMV and TMV, and PZSV, TAV and TuYV, respectively. In the environmental effect network (**Figure 3B**), generated from the subtraction of the CRF-MEMs matrix from that of the CRF-Full model, the set of interactions associated with virus taxonomy and/or mode of transmission (BChV, BMYV and TuYV; PMoV and PZSV; and PMMoV and TMGMV) were maintained, as well as the role of hub by CMV and PMoV, which interacted with PMMoV, TAV, TCV, and TMV, and with PZSV, TAV and TuYV, respectively. Last, the habitat effect network (**Figure 3C**), generated by subtracting the matrix of the CRF-Host model from that of the CRF-Full model, revealed again the interactions associated with shared evolutionary history and/or mode of transmission (BChV, BMYV, TuYV; PMoV, PZSV; PMMoV, TMGMV), but showed a diminished role of CMV and PMoV as hubs structuring the network, indicating that some of the positive interaction involving these two viruses (i.e., CMV and PMMoV, CMV and WMV, PMoV and TAV) were due to host-related structuring of infections.

**Figure 3.**
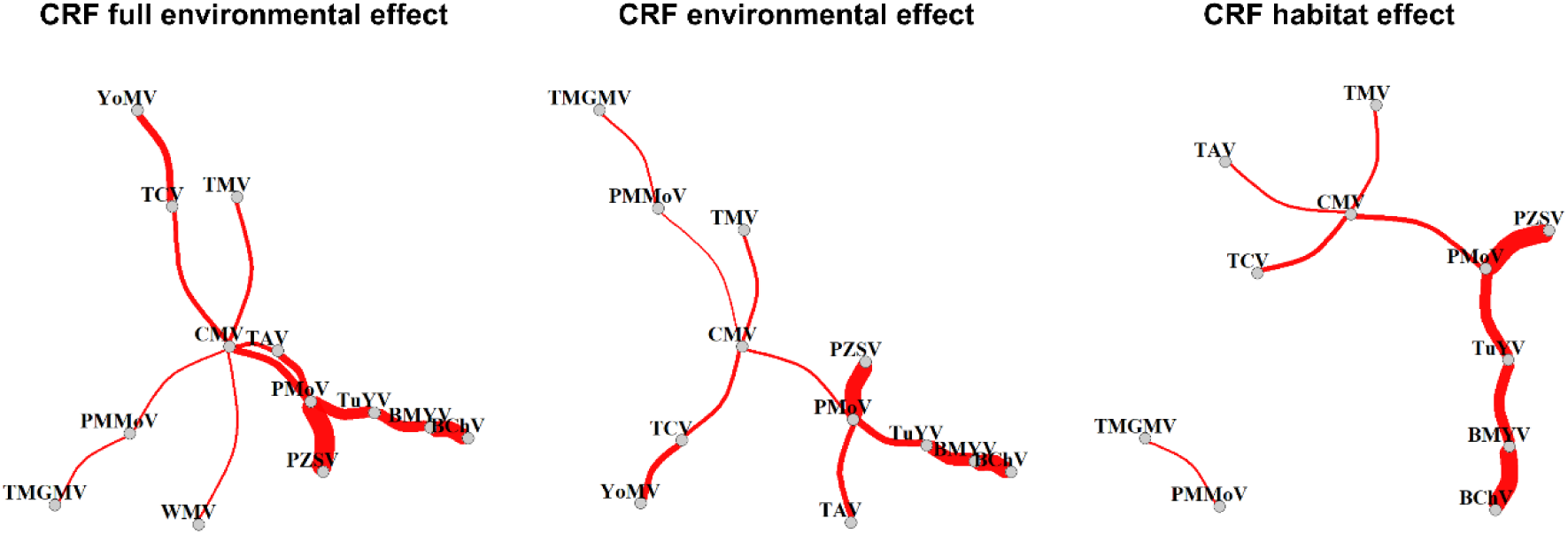
Virus interaction networks. A) Conditional Markov random field interaction network accounting for all spatial, environmental and host effects (CRF-Full). B) Conditional Markov random field interaction network accounting for all host and environmental effects. C) Conditional Markov random field interaction network accounting for the habitat effects. **I**nteractions with an absolute edge weight ≤ 0.05 are not shown in the networks. Alt text. Associations between pairs of viruses after accounting for environmental variables.

Thus, a set of interactions that involved viruses of shared evolutionary history and/or mode of transmission (BChV, BMYV, TuYV; PMoV, PZSV; PMMoV, TMGMV) were present in the networks derived from all models (**Figures 2** and **3**).

## DISCUSSION

The current understanding of mechanisms that determine virus interactions in coinfected plants (7, 20), is limited by the lack of information available on plant virus ecology and the structuring of viral communities (18, 19). Here we elucidate on factors that structure virus coinfection in plant communities of four habitats that characterise an ecosystem of central Spain. Analyses showed that more than half (∼ 55%) of the 1,931 individuals of 94 plant species were coinfected by two or more virus species. This prevalence of coinfection by the 18 virus species in this study is similar to that of another performed 15 years earlier in the same geographic region that involved 11 virus species (28). Studies of coinfection in crops and wild plants (11–13, 15–17, 19, 40) show estimates of coinfection prevalence that range from 5.6% (four viruses in wild sweet potato relatives in Uganda) to 70% (five viruses in California wild grasslands). The relationships reported between coinfection prevalence and host or virus species is unclear and suggests that coinfection depends on a combination of factors intrinsic to the coinfecting viruses, and/or on extrinsic ones related to the host and environment. In our study, both the frequency of coinfection and the number of coinfecting viruses depended largely on habitat, being more prevalent in Crop and Edge than in Wasteland and Oak. Additionally, Edge had a major role in structuring coinfection, consistent with previous work (28). The results suggest extrinsic factors drive non-random virus associations through two main processes, a common response to ecological heterogeneity, and indirect associations among subsets of viruses (4, 18, 41). Our findings also showed that coinfection was not solely a function of extrinsic determinants.

We analysed the potential role of extrinsic factors using random fields models (18). Analyses of the correlation between the infection distributions of each pair of viruses showed 66 associations that involved all virus species except BYDV (**Figure S5**). The MRF network revealed 15 significant interactions. Thus, most associations detected by correlation analyses of infection distributions were due to indirect interactions derived from multispecies distributions, a result also reported by (18), which supports the first null hypothesis (the proportion of random and non-random associations is equivocal). Interestingly, none of the CRF models conditioned on environmental variables, outperformed the MRF one. Only the models that included the spatial (CRF-MEMs) and host-related (CRF-Host) variables were similar to MRF model in terms of AUC and predictive performance (sensitivity and specificity). The virus interactions retained in these CRF models were subsets of those of the MRF model, with a hub of interactions common to the three models with virus pairs that share a mode of transmission (BChV, BMYV and TuYV; PMoV and PZSV; PMMoV and TMGMV), plus interactions of CMV and PMoV with viruses with other evolutionary histories and/or with different modes of transmission that varied among models (**Figure 2**). However, when the MRF adjacency matrix was subtracted from the CRF-Full matrix, which included all spatial and environmental variables in **Table 2**, all the positive interactions of the MRF network were retained (compare **Figure 2A** and **Figure 4A**), indicating that a common response by viruses to the environmental conditions determined this set of coinfections. The subtraction of the adjacency matrix of host variables from that with all environmental variables resulted in the loss of interactions between CMV and PMoV, although the hubs of interactions above were retained (**Figure 3C**), indicating these coinfections did not depend on host variables. These results do not support our second null hypothesis (extrinsic factors have no effect on the proportion of random to non-random associations) but show that common responses to environmental heterogeneity play a major role in determining non-random associations among the 18 virus species in our data set. This finding is consistent with results from studies of virus infection in grasses in California (19) and in *Plantago lanceola*ta in Finland (18). The conditional probability models indicated that a common mode of transmission is a driver of coinfection, as shown by consistent associations between the contact-transmitted TMGMV and PMMoV, between viruses transmitted by aphids in a persistent manner (TuYV, BMMV, BChV), and between the two pollen transmitted viruses PZSV and PMoV. These results suggest that the virus trait responses related to the mode of transmission are central to coinfection. Co-transmission of parasites that share a transmission mode has been reported as a driver of multiple infections (4). In plant viruses, it has been shown that viruses transmitted by the same vector are more likely to coinfect (14), and host plants preferred by foraging vectors have a higher prevalence of coinfections (19, 42). Both processes may explain the non-random association in coinfected plants of the three poleroviruses that had a similar host range, BMYV, BChV and TuYV (McLeish et al., 2024), as persistently transmitted viruses have high vector specificity, and aphid vectors may also have a high host plant preference (43, 44). Similarly, host preference by insect pollinators (45, 46) may favour the coinfection of pollen transmitted viruses and of tobamoviruses, which may be transmitted non-specifically by pollinators (47, 48). Conversely, there were few associations, not consistent among models, between viruses non-persistently transmitted by aphids, which show little vector specificity and are not transmitted preferentially to aphid-preferred plants during aphid feeding (44). Interestingly, only one non-random association, detected in only one model (CMV and WMV, in MRF) involved a potyvirus partner. As potyviruses have been shown to establish synergistic interactions with viruses from other genera in many different hosts (20), this result argues against synergism being a major driver of virus associations. Within-host processes also seemed to have a minor role in coinfections of barley/cereal dwarf viruses (14).

Our study is a considerable addition to the understanding of drivers of plant virus coinfection. We show a high frequency of coinfection in our diverse multi-host – multi-virus system. We also show that only a minor fraction of coinfections are non-random associations. The large fraction of random associations in coinfected plants supports hypothesis I, and is consistent with ecological fitting, which our earlier work has shown to play a major role in virus infection distributions in the analysed ecosystem (49–51). Consistent with other reports, our results underscore the relevance of common responses to ecological heterogeneity driving coinfection, and strongly suggests that the mode of horizontal transmission plays a role in these responses. Data suggest that direct virus interactions in the coinfected hosts have a minor role in non-random associations. Rather, results highlighted a major role of between-host, rather than within-host processes in driving coinfection. Assessing how general this pattern is will require further studies.

## ACKNOWLEDGEMENTS

We thank Antolín López Quirós and Miguel Ángel Mora for technical support.

## DATA AVAILABILITY

All data are incorporated into the article and its on line supplementary material.

## AUTHOR CONTRIBUTIONS

A.D.Z. and B.M.B. obtained the data; A.D.Z. analysed the data; M.J.M. and F.G.- A. designed the study; A.D.Z., M.J.M. and F.G.-A. wrote the initial version of the manuscript, and all authors contributed to the final one.

